# Dauer diapause has transgenerational effects on starvation survival and gene expression plasticity

**DOI:** 10.1101/342113

**Authors:** Amy K. Webster, James M. Jordan, Jonathan D. Hibshman, Rojin Chitrakar, L. Ryan Baugh

## Abstract

Phenotypic plasticity is facilitated by epigenetic regulation, and remnants of such regulation may persist after plasticity-inducing cues are gone. However, the relationship between plasticity and transgenerational epigenetic memory is not understood. Dauer diapause in *Caenorhabditis elegans* provides an opportunity to determine how a plastic response to the early-life environment affects traits later in life and in subsequent generations. We report that after extended diapause, post-dauer worms initially exhibit reduced reproductive success and greater inter-individual variation. In contrast, F3 progeny of post-dauers display increased starvation resistance and lifespan, revealing potentially adaptive transgenerational effects. Transgenerational effects are dependent on the duration of diapause, indicating an effect of extended starvation. In agreement, RNA-seq demonstrates a transgenerational effect on nutrient-responsive genes. Further, post-dauer F3 progeny exhibit reduced gene expression plasticity, suggesting a trade-off between plasticity and epigenetic memory. This work reveals complex effects of nutrient stress over different time scales in an animal that evolved to thrive in feast and famine.

---

Epigenetic regulation mediates phenotypic plasticity, such that a single genotype can produce different phenotypes in response to environmental conditions (1). Most epigenetic modifications are reset at the beginning of each generation, but those that persist have the potential to impact environmental adaptation and evolution independent of DNA sequence (2-4). Consequently, there is substantial interest in identifying environmental perturbations that elicit transgenerational epigenetic effects. Epigenetic inheritance has been reported in the roundworm *Caenorhabditis elegans*, but these studies typically focus on an aphysiological stimulus, such as a mutation or exogenous RNAi trigger (5, 6), or they assay a molecular rather than organismal phenotype, such as gene expression (7-10). Consequently, the importance of epigenetic inheritance to environmental adaptation is unclear. Identification and characterization of plasticity-induced transgenerational effects manifest at the organismal level can address important questions. Do epigenetic changes in gene regulation affect organismal traits? Does epigenetic memory provide a predictive adaptive response that improves fitness? Alternatively, does epigenetic memory arise due to a failure to reset the epigenetic program between generations, potentially posing a fitness cost?

*C*. *elegans* has a variety of developmental responses to nutrient availability (11-14). In particular, larvae respond to specific environmental cues (high population density, low food availability, and high temperature) and undergo an alternative developmental program resulting in dauer arrest (15). Dauer arrest is a classic example of diapause since dauer larvae develop in response to cues perceived in advance of arrest (16). Dramatic morphological and gene regulatory changes occur during dauer development (17-19) under the regulation of highly conserved signaling pathways (20-24). Dauers have a thicker cuticle and plugged pharynx, supporting increased stress resistance but preventing feeding. Dauers also have an altered nervous system, gut, and muscles compared to non-dauers (25-27). They are provisioned with fat stores that aid survival during prolonged starvation (28, 29). If environmental conditions improve, larvae exit dauer arrest and ultimately become reproductive adults.

*C. elegans* collected from the wild are mostly in the dauer stage, providing evidence that starvation stress during dauer diapause is a common feature of their life history (30). Time spent in dauer does not alter the length of the adult lifespan, suggesting an “ageless” state (31). Nonetheless, dauer larvae do eventually die, with about one-third failing to recover at 60 days (31). Further, germline defects increase in frequency as the duration of dauer diapause increases (32), suggesting that at least some consequences of starvation incurred during diapause persist in post-dauers. It is currently unknown if effects of long-term dauer diapause persist upon recovery to affect important life-history traits, including progeny quality and starvation resistance. The consequences of long-term dauer diapause beyond one generation are, to our knowledge, completely unexplored.

Here, we report the consequences of long-term dauer diapause over multiple generations. We found that long-term dauers recovered to produce fewer, smaller and starvation-sensitive F1 progeny that exhibited greater inter-individual variation, suggesting proximal fitness costs. In contrast, great-grandprogeny (F3 generation) of long-term dauers exhibited increased starvation resistance and lifespan, consistent with potentially adaptive transgenerational effects. Increased starvation resistance was dependent on the time spent in the ancestral dauer diapause, suggesting that it is starvation during dauer, rather than dauer development itself, leading to these transgenerational effects. Consistent with this, we identified epigenetic differences in nutrient-responsive gene expression in the F3 generation including a dampened response to nutrient availability. These results indicate that ancestral environment can influence how worms respond to their current environment, and they suggest a trade-off between epigenetic memory and plasticity.

## RESULTS

We carefully controlled worm population density and bacterial food concentration in a liquid culture system to cause wild--type worms to either enter dauer diapause or to bypass it (18) (Fig 1). Ancestors of dauer and control worms were well fed for multiple generations (>3) to control for multigenerational effects of starvation or dietary restriction. Long-term dauers remained dauers for 36-45 days, and then were put on plates with food to recover. In parallel, control worms that did not experience dauer were plated from liquid culture so that post--dauer and control worms were paired for experiments.

**Fig 1:**
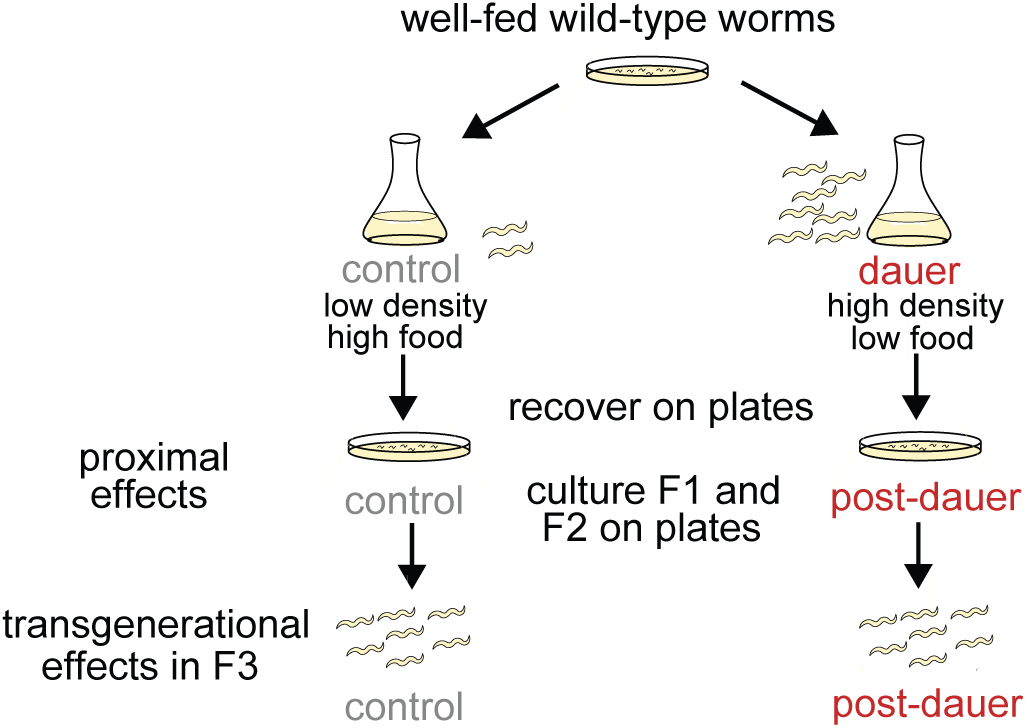
Schematic of experimental design. Well-fed, wild-type worms were hypochlorite treated as gravid adults to obtain embryos, which were placed in dauer or control conditions. For long-term dauer conditions, worms remained in culture for 40-49 days and were dauers for 36-45 days. Worms from dauer and control cultures were plated on food in parallel. These lay F1 progeny, some of which were used for experiments of proximal effects of long-term dauer diapause. Other F1 adults were hypochlorite treated to obtain F2 embryos. F2 adults were hypochlorite treated to obtain F3 embryos. F3 post-dauer and control worms were used for experiments.

### Long-term dauer diapause reduces reproductive success in the P0 generation

We determined the proximal consequences of long--term dauer diapause on reproductive success. Following diapause, post-dauer worms recovered to produce fewer F1 progeny (Fig 2A and Supplementary Fig 1A). Post-dauer adults also produced smaller embryos (Fig 2B and Supplementary Fig 1B). *C. elegans* embryos hatch as L1 larvae. In the absence of food, L1 larvae survive starvation by remaining arrested as L1s, and this is reversible upon feeding. Survival during L1 arrest is a measure of starvation resistance (12). We found that post--dauers produced F1 progeny that were relatively sensitive to starvation by two metrics: L1 starvation survival (Fig 2C) and growth rate during recovery from L1 starvation (Fig 2D and Supplementary Fig 1C). Thus, long-term dauer arrest incurs costs to average fecundity as well as to progeny quality and starvation resistance, indicating reduced reproductive success.

**Fig 2:**
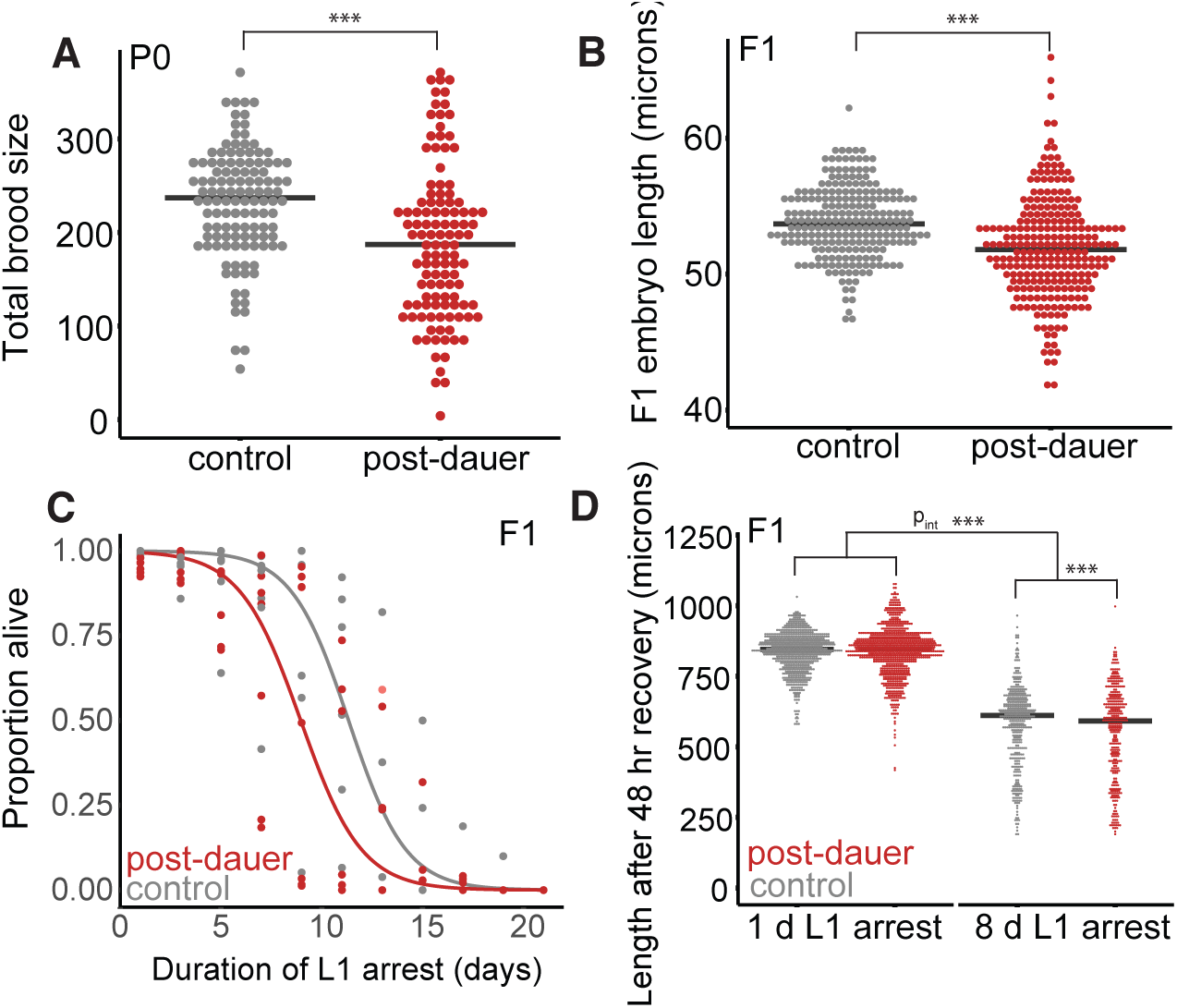
Long-term dauers recover to produce fewer, smaller F1 progeny that are starvation sensitive. A. Brood sizes following long-term dauer diapause or control conditions. 7 biological replicates were scored with 13-18 individuals per biological replicate per condition. Effect of condition, p = 4.46 x 10^−7^. B. Embryo length in the F1 progeny of post-dauers and controls. 6 biological replicates were scored with 23-70 individual embryos scored per biological replicate. Effect of condition, p = 9.0 x 10^−13^. C. L1 starvation survival for 7 biological replicates of F1 progeny of post-dauers and controls. Logistic regression curves were fit, and median survival was determined for each replicate. Paired t--test on median survival, p = 0.10. D. Worm length after 48 hr of recovery from 1 or 8 days of L1 arrest was scored in the F1 progeny of postdauers and controls. 8 biological replicates were scored. For 1 day, 50-335 worms were scored per biological replicate; following 8 days, 7-118 worms scored per replicate. Effect of condition for 1 day, p = 0.95; effect of condition for 8 days, p < 2.0 x 10^−16^. A linear mixed-effect model was fit with condition and length of starvation as fixed effects and biological replicate as a random effect. An interaction term was included for fixed effects. Effect of the interaction of condition and length of starvation, p = 5.1 x 10^−14^; effect of starvation, p < 2.0 x 10^−16^; effect of condition, p = 0.20. A,B,D. Linear mixed--effect models were fit with condition (post--dauer vs. control) as a fixed effect and biological replicate as a random effect. P-values were calculated using the Wald test. Horizontal black lines represent medians. For means of biological replicates and grand mean, see Supplementary Fig 1. ***p<0.001

Because we identified differences in *mean* trait values for post-dauer and control worms upon recovery and in the F1 generation, we also considered the possibility that experiencing long-term dauer diapause could alter the inter-individual *variation* of key traits within a worm population. We found that post-dauer worms exhibited greater variation in brood size than controls, and that F1 embryos were more variable in length (Supplementary Fig 2A-B). Consistent with these observations, once these embryos hatched as L1s, they appeared to exhibit greater variation in body length after recovery from 8 days of starvation, but not 1 day of starvation (Supplementary Fig 2C). This suggests that increased inter-individual variation is uncovered by the stress of extended L1 starvation. Together our results suggest that the proximal consequences of long-term diapause include decreased reproductive success and increased phenotypic variation.

### Long-term dauer diapause increases starvation resistance and lifespan transgenerationally

We asked whether effects of long-term dauer diapause persisted beyond the F1 generation to the F3 generation. Persistence to the F3 generation suggests transgenerational epigenetic inheritance, while effects in the F1 and F2 generations could be due to maternal effects (33). We measured starvation resistance in three ways: L1 starvation survival, body length following 48 hours of recovery from L1 starvation, and brood size following recovery from L1 starvation. In contrast to F1 L1 larvae, we found that F3 progeny of long--term dauers exhibited increased L1 starvation survival (Fig 3A). Likewise, F3 progeny of long--term dauers allowed to recover for 48 hr from 8 days of L1 starvation were larger than controls recovering from 8 days of L1 starvation, demonstrating increased growth rate (Fig 3B and Supplementary Fig 3A). In addition, the broods of F3 progeny of long-term dauers following 8 days of L1 starvation were larger than the broods of F3 controls following 8 days of L1 starvation (Fig 3C and Supplementary Fig 3B). We also assayed growth rate and brood size in the F3 post-dauers with just 1 day of L1 starvation and found no significant differences in these traits (Fig 3B-C and Supplementary Fig 3A-B). In addition, we did not find evidence of differences in inter-individual variation in the F3 generation (Supplementary Fig 3F-G), suggesting that the differences in mean trait values in the F3 generation were not due to a difference in the shape of the underlying trait distribution. Together these data support the conclusion that long-term dauer diapause leads to epigenetic inheritance of increased starvation resistance in F3 progeny.

**Fig 3:**
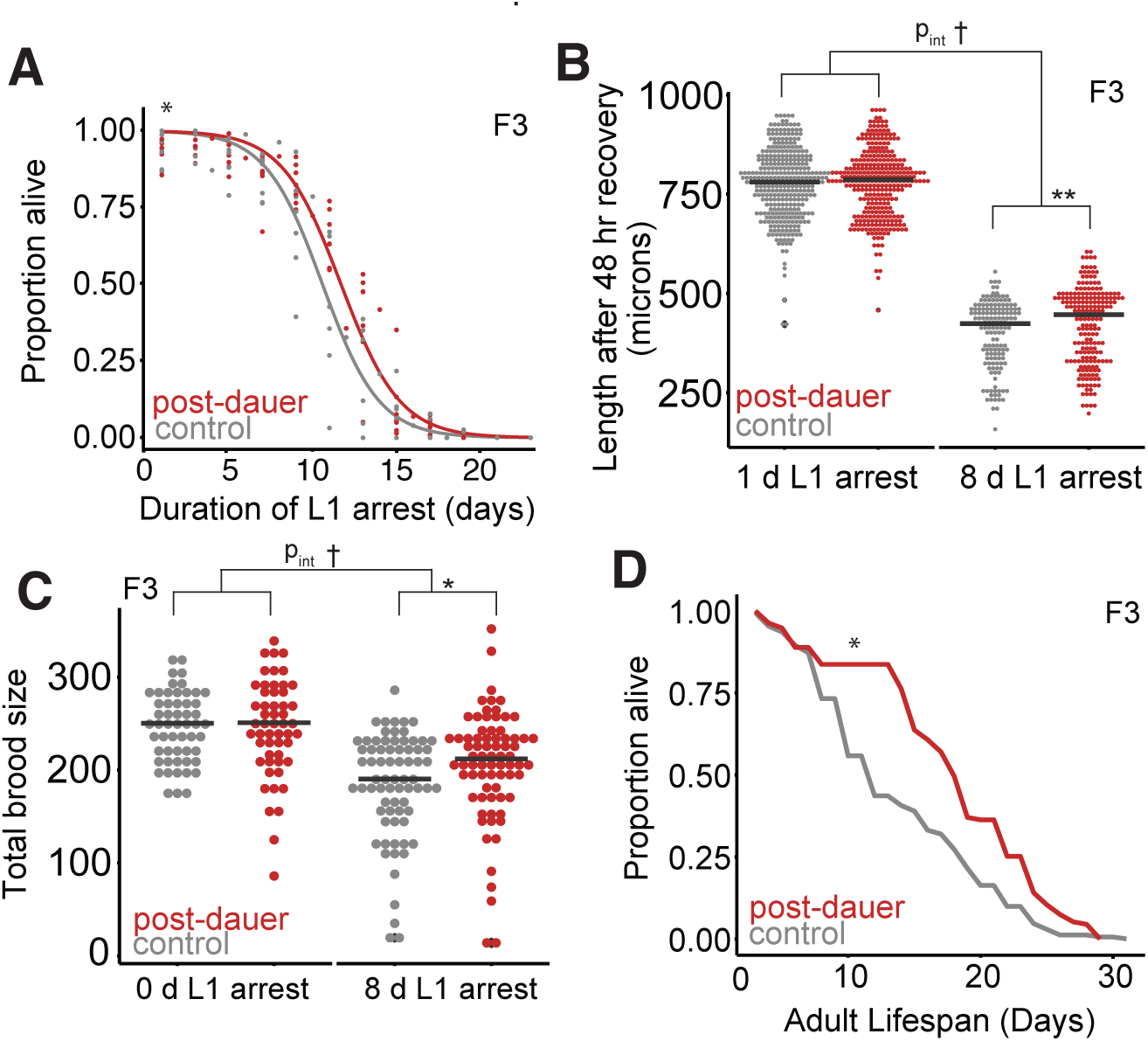
F3 progeny of long-term dauers exhibit increased starvation resistance and lifespan. A. L1 starvation survival was scored in the F3 progeny of post-dauers and controls. 8 biological replicates were scored, logistic curves were fit, and median survival times were determined. Paired t-test on median survival, p=0.01. B. Worm body length following 48 hours of recovery from either 1 or 8 days of L1 arrest. 4 biological replicates were scored, consisting of 342 controls starved 1 day, 323 post-dauers starved 1 day, 148 controls starved 8 days, and 204 post-dauers starved 8 days. Effect of condition for 1 day, p = 0.25; effect of condition for 8 days, p = 0.0048. Effect of the interaction between condition and length of starvation, p = 0.066; effect of the length of starvation, p < 2.0 x 10^−16^; effect of condition, p = 0.93. C. Brood size was scored for F3 progeny of controls and post-dauers that experienced 0 or 8 days of L1 arrest. 3 biological replicates for 0 days; 5 biological replicates for 8 days. For 0 d L1 arrest, 54 controls and 53 post-dauers; for 8 d of L1 arrest, 72 controls and 74 post-dauers. Effect of condition for 0 days, p = 0.82; effect of condition for 8 days, p = 0.03. Effect of the interaction between condition and length of starvation, p = 0.087; effect of the length of starvation, p < 1.0 x 10^−4^; effect of condition, p = 0.87. D. Lifespans of at least 135 F3 progeny of post-dauers and controls from 3 biological replicates (See Supplementary Fig 3C-E for individual replicates). Paired t-test on the means of biological replicates, p = 0.026. B,C. Linear mixed-effect models were fit with condition (post-dauer vs. control) as a fixed effect and biological replicate as a random effect. Next, a linear mixed-effect model was fit with condition (postdauer vs. control) and length of starvation (0 or 1 vs. 8 days) as fixed effects and biological replicate as a random effect. An interaction term was included for fixed effects. P-values were calculated using the Wald test. *p<0.05, **p<0.01, ***p<0.001; t interaction p<0.1; n.s. not significant. Horizontal black lines represent medians. For means of biological replicates, see Supplementary Fig 2.

We found transgenerational effects of long-term dauer arrest that were present beyond the L1 stage and into adulthood in F3 worms that were never starved. F3 progeny of long-term dauers lived longer as adults than controls (Fig 4D, Supplementary Fig 3C-E), demonstrating that there are transgenerational consequences of long-term dauer diapause outside the context of starvation.

**Fig 4:**
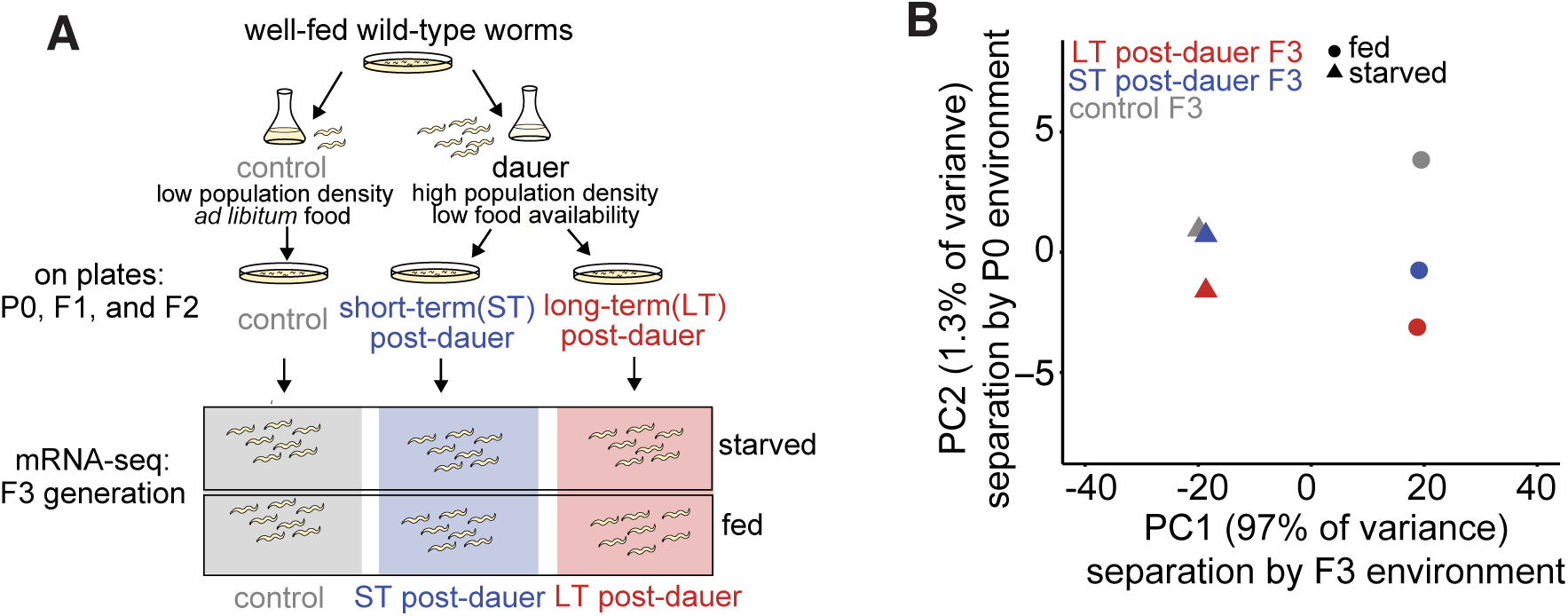
mRNA-seq reveals relative contributions of current environment and ancestral environment in shaping gene expression variation. A. Schematic of experimental set-up for collection of RNA-seq samples. B. Principal component analysis (PCA) of 6 conditions including all 8649 reliably detected genes. 97% of variance explained by whether worms were fed or starved at collection (PC1). 1.3% of variance explained by whether ancestors experienced control, short-term dauer, or long-term dauer conditions (PC2).

We next wanted to determine whether the transgenerational increase in starvation resistance depends on the duration of dauer diapause. When worms were dauers for just six days, F3 progeny of these short--term dauers did not exhibit differences in starvation survival or brood size following starvation (Supplementary Fig 4). We had the statistical power to detect differences in starvation survival of less than a day (Supplementary Table 1; power analysis not shown). This suggests that the transgenerational effects of experiencing long-term dauer arrest are dependent on the length of starvation during arrest as opposed to the dauer-inducing culture conditions, dauer formation, or dauer recovery.

### Epigenetic memory of dauer diapause affects gene expression globally

Transgenerational effects of long-term dauer diapause on important organismal traits suggest epigenetic regulation of gene expression. We used mRNA-seq to determine if gene expression patterns in both fed and starved F3 progeny of controls, short-term dauers, and long-term dauers differed (Fig 4A). This two-factor design allowed us to analyze both transgenerational and instantaneous effects of nutrient availability on gene expression. We performed principal component analysis (PCA) on the normalized mean counts per million (CPM) values for six conditions as well as on individual biological replicates (Fig 4B, Supplementary Fig 5). PC1 separated condition means depending on whether the worms were fed or starved as F3 L1s, while PC2 separated them according to length of time the initial generation spent in dauer diapause. Notably, instantaneous effects of nutrient availability (PC1) appeared to explain substantially greater variance in gene expression than epigenetic effects (PC2). Nonetheless, these results suggest that ancestral environmental conditions play a small, but detectable, role in shaping gene expression in the F3 generation.

We compared gene expression plasticity in F3 long-term post-dauers to controls. That is, we assessed gene expression differences between starved and fed F3 worms with long-term dauer ancestors to starved and fed F3 worms with control ancestors (Fig 5A). Regardless of ancestral background, starved and fed worms exhibited dramatic gene expression differences (Fig 5A, Supplementary File 1), consistent with PC1. Gene expression responses to nutrient availability were also highly correlated. Notably, virtually no genes had large differences in one comparison and not the other. However, the slope of the linear regression was significantly less than 1, suggesting that gene expression plasticity in response to nutrient availability in the F3 generation following long-term dauer diapause is dampened relative to controls.

**Fig 5:**
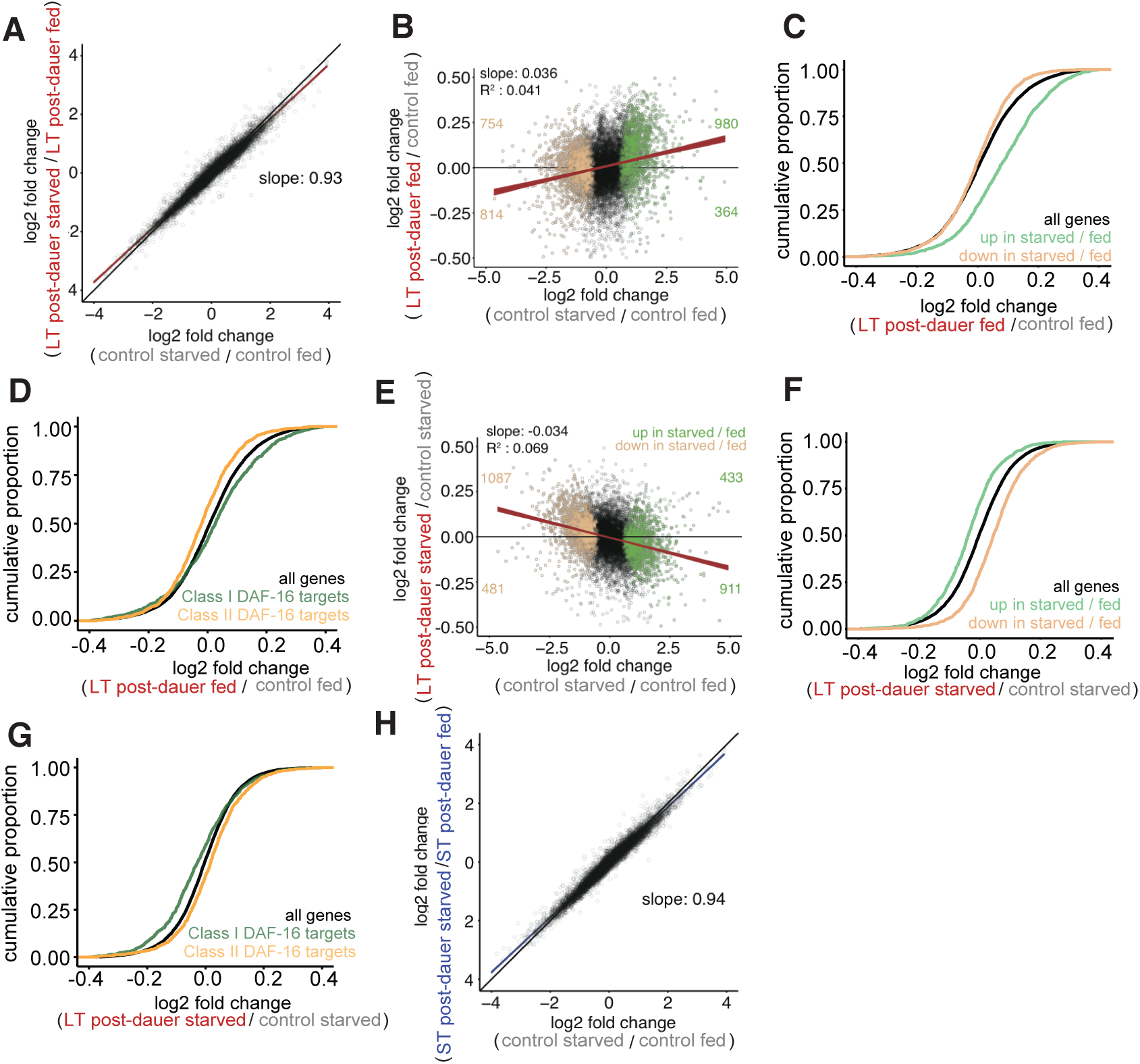
F3 progeny of dauers exhibit reduced gene expression plasticity driven by differences in both fed and starved worms. A. Log2 fold changes of all genes in control starved / control fed plotted against LT post-dauer starved / LT post-dauer fed. Red line is linear regression through all points and width indicates 95% confidence interval (CI). Black line is the line y = x. Slope is 0.93, with CI from 0.926 to 0.934. B. Log2 fold changes of all genes in control starved / control fed plotted against log2 fold changes of LT postdauer fed F3 / control fed plotted in black. Slope is significantly different from 0, p < 2.2 x 10^−16^ C. Genes up in starved / fed: p = 0; to genes down in starved / fed: p = 3.3 x 10^−8^. D. Class I and Class II targets defined in Tepper et al. Class I targets: p = 1.6 x 10^−5^; to Class II targets: p = 2.3 x 10^−11^. E. Log2 fold changes of all genes in control starved / control fed plotted against log2 fold changes of LT post-dauer starved F3 / control starved plotted in black. Slope is significantly different from 0, p < 2.2 x 10^−16^. F. Genes up in starved / fed: p = 0; to genes down in starved / fed: p = 0 G. Class I targets: p = 1.0 x 10^−12^; to Class II targets: p = 4.0 x 10^−7^. H. Log2 fold change of all genes in control starved / control fed compared to log2 fold change in ST post-dauer starved / ST post-dauer fed. Blue line indicates simple linear regression through all points, and thickness of line indicates 95% CI. Slope is 0.94, with CI from 0.936 to 0.943. B,E. Genes up in control starved / control fed using FDR 1 x 10^−10^ are plotted in green. Genes down in control starved / control fed are plotted in tan. Red line is a simple linear regression through all points, and thickness of line indicates 95% confidence interval. Number of genes differentially expressed in control starved / control fed in each quadrant is indicated. C,D,F,G. Within the indicated comparison, cumulative distribution functions (CDFs) of indicated gene lists are plotted. Kolmogorov-Smirnov test used to assess significance of gene list distribution compared to all genes.

Dampened plasticity could occur if fed F3 progeny of long-term dauers had a transcriptional profile that appeared “less fed” compared to fed controls or if starved F3 progeny of long-term dauers appeared transcriptionally “less starved” compared to starved controls. First, we found fed F3 progeny of long-term dauers appeared less fed (or, equivalently, more starved) relative to fed controls by comparing the effect of plasticity in controls to the epigenetic effect in fed larvae (Fig 5B,C). We further tested this effect using the most differentially expressed nutrient-responsive genes. These genes, highlighted in Fig 5B, showed that genes that were significantly up- or down-regulated in starved worms compared to fed worms tend to also be up- or down-regulated, respectively, in fed F3 progeny of long--term dauers relative to fed controls (Figs 5B,C). We also found that the regulatory targets of the transcription factor *daf-16* (34), which promotes starvation survival (12), overlap with the nutrient-responsive genes and display a similar epigenetic effect of long-term dauer diapause (Fig 5D, Supplementary Fig 6A). This result corroborates the epigenetic effect on nutrient-responsive genes using gene lists defined outside the context of this study. These results provide evidence that dampened gene expression plasticity is driven at least in part by differences in nutrient--responsive gene expression in fed F3 progeny of long-term dauers.

We wanted to determine whether there are differences in nutrient-responsive gene expression in the *starved* F3 progeny of long-term dauers compared to controls. In contrast to the fed F3 progeny, the starved F3 progeny of long-term dauers exhibited a negative correlation with the control nutrient response (Fig 5E,F). This was corroborated by the fact that *daf-16* target genes displayed similar behavior (Fig 5G). Together these results suggest that there are transgenerational effects on gene expression in both fed and starved F3 larvae that collectively contribute to reduced gene expression plasticity.

F3 progeny of short-term dauers did not display detectable epigenetic effects on starvation resistance, but we did detect effects on gene expression. We found that F3 progeny of short-term dauers showed similar transcriptional changes as identified in the F3 progeny of long-term dauers, including dampening of nutrient-responsive gene expression (Fig 5H, Supplementary Fig 6B-D). However, transgenerational changes in gene expression following short-term dauer appear to be less consistent and possibly of a smaller magnitude than those elicited by long-term dauer. PC2 aligns means of short-term dauer samples between control and long-term dauer samples (Fig 4A). The slope of the dampening response of F3 progeny of short-term dauers compared to controls was closer to 1 than F3 progeny of long-term dauers, and F3 progeny of long-term dauers exhibit dampening relative to short-term dauers (Fig 5H, Supplementary Fig 6B). In addition, we examined gene expression shifts within individual paired replicates (replicates in which an F3 dauer sample was collected in parallel with a control). All paired replicates for starved F3 progeny of long-term dauers compared to starved controls showed shifts in nutrient-responsive genes in the expected direction (Supplementary Fig 6E). Paired replicates for F3 progeny of short-term dauers exhibited shifts in gene expression for the majority, but not all, pairs (Supplementary Fig 6F). Collectively, these results suggest that transgenerational epigenetic effects of short-term dauer are detectable and qualitatively similar to those of long-term dauer, but that they are less consistent, likely accounting for lack of detectable effects on starvation resistance.

## DISCUSSION

We utilized a well-studied model of phenotypic plasticity, dauer diapause, to interrogate the proximal and transgenerational consequences of early-life environment on organismal traits and gene expression. Post-dauer worms displayed reduced reproductive success after long-term diapause, but their F3 progeny exhibited increased starvation resistance and lifespan. F3 progeny of both short- and long-term dauers exhibited changes in gene expression in nutrient-responsive genes, consistent with a reduction in gene expression plasticity in response to nutrient availability. This work has broad implications for understanding the consequences of starvation, including the potential for epigenetic inheritance in response to nutrient availability and potentially adaptive responses across generations.

Long-term dauers recovered to produce fewer, smaller F1 progeny that are starvation sensitive compared to worms that do not enter dauer. We also found that long-term post-dauers exhibit increased inter-individual variation, consistent with developmental decanalization (35) stemming from deleterious effects of extended starvation (See Supplementary Discussion). Dauer diapause is evolutionarily adaptive, as it allows worms to survive conditions that would otherwise be lethal. Our results suggest that while it is adaptive to be able to enter dauer to survive adverse conditions, there is a cost to extended dauer diapause. The proximal costs of long-term dauer diapause are consistent with our lab’s previous work identifying costs to eight days of L1 starvation, as worms recovered from L1 starvation are more sensitive to subsequent starvation and produce smaller broods with decreased progeny quality (36). Both extended L1 starvation and dauer diapause appear costly, but this is likely due to the costs of starvation experienced during these states, rather than simply arresting development. Post-dauer worms that experienced dauer for just one day live longer and produce larger broods (37). Together these observations suggest that hormesis, in which a mild stress can increase fitness (38), may occur when starvation is brief, but that extended starvation is costly as the buffering capacity of the organism is overwhelmed.

We found that the F3 progeny of long-term dauers exhibit increased starvation resistance as larvae and live longer as fed adults, indicating that transgenerational effects manifest in both fed and starved animals and that they persist through the life cycle. Multigenerational consequences of L1 starvation have been previously examined and exhibit some similarities to those of long-term dauer diapause. Following 8 days of L1 starvation, we previously observed increased starvation resistance in the F2 but not F3 generation (36). We did not detect a reproducible effect on lifespan following L1 starvation in any generation, though increased lifespan in the F3 generation following L1 starvation has been reported in another study (39). Notably, transmission of increased starvation resistance to F1 and F2 progeny of worms that experienced L1 starvation required sorting larvae by size after 2 days of recovery from starvation (36), and in the present study such sorting was not required. Still, both L1 and dauer starvation paradigms reveal apparent proximal fitness costs of extended starvation followed by potentially adaptive increases in starvation resistance in subsequent generations. The switch from proximal starvation sensitivity to heritable starvation resistance in both paradigms is of particular interest.

In addition to life-history trait differences, we identified transgenerational effects of dauer diapause on gene expression plasticity in response to nutrient availability. Specifically, we found that F3 progeny of both short-term and long-term dauers exhibited reduced gene expression plasticity, with starved F3 worms appearing less starved and fed F3 worms appearing less fed. Theory suggests that there should be costs and limits to plasticity, though there are relatively few empirical examples of such costs (40, 41). While we emphasize that the transgenerational effect on gene expression plasticity does not necessarily indicate plasticity at other levels of regulation, one interpretation of this result is that maintaining an epigenetic memory of ancestral environmental conditions limits plastic responses to current environmental conditions. In other words, there is hypothetically a fixed capacity to respond to environmental conditions, but this capacity can be exhausted by instantaneous and epigenetic regulation.

It is possible that subtle changes in starvation resistance and gene expression reflect physiological fine-tuning in anticipation of future conditions based on ancestral history. In contrast, it is possible that epigenetic memory of ancestral conditions occurs but is not necessarily adaptive, as a potentially neutral or even costly by-product of some of other selected trait. Epigenetic stability from parent to offspring is favorable given that four conditions are met: 1) the environment is variable; 2) parental environment has some predictive power over offspring environment; 3) transgenerational effects increase fitness of parents and/or offspring; and 4) costs associated with the transgenerational response are low (42). Though it is clear that *C. elegans* experience variable nutrient conditions in the wild (43), it is currently unknown how predictive parental environment is of offspring environment over multiple generations. Here, we find that F3 progeny of long-term dauers are more starvation resistant, and this is not accompanied by detectable costs in growth or fecundity, consistent with the possibility that this tra nsge ne rational effect is adaptive. Recognizing that it is ultimately difficult to determine if such a complex trait as epigenetic inheritance is evolutionarily adaptive, we hope that future studies in the context of the natural ecology of *C. elegans* will shed light on this question.

## MATERIALS AND METHODS

### Dauer and control cultures

The wild-type strain N2 was used for all experiments. N2 was obtained from the Sternberg collection at the California Institute of Technology, originally from the CGC in 1987. Worms were maintained for at least 3 generations in standard laboratory conditions without starving prior to beginning experiments. ~10 adults were picked onto each of 4-5 10 cm NGM plates seeded with *E. coli* OP50 and maintained at 20°C. Embryos were obtained by standard hypochlorite treatment after four days in culture. For dauer-forming conditions, embryos were suspended in S-complete at a density of 5 per microliter with 1 mg/mL *E. coli* HB101(18). HB101 was prepared as described previously (44). For control conditions, embryos were suspended at 1 per microliter with 38 mg/ml HB101. Worms were cultured at 180 rpm and 20°C in 25 mL glass Erlenmeyer flasks in a volume of 5 mL (25,000 worms for dauer conditions and 5,000 for control conditions). For experiments that required greater than 5,000 control worms, a 20 mL volume in a 250 mL Erlenmeyer flask with the same density and food concentration was used. Dauer formation occurs in approximately 4 days with nearly 100% penetrance. Short-term dauers remained in culture for 10 days, being arrested as dauers for 6 days. Long-term dauers remained in culture for 40-49 days, being arrested as dauers for 36-45 days. Survival was 90 - 100% in long-term dauer cultures. Control worms were in culture for 40-44 hours and were plated as L4 larvae. The L4 stage was chosen because dauers recover to become L4 larvae.

### Dauer recovery and maintenance

Dauer and control conditions were paired such that control cultures were set up to be recovered at the same time as the dauer culture. Thus, recovery, maintenance, sampling and assaying were done in parallel. Worms were taken from liquid cultures and plated on 10 cm NGM plates seeded with OP50 and incubated at 20°C. To obtain F1 progeny, approximately 1000 P0 worms were plated per seeded plate, and these were hypochlorite treated 2 days later to obtain F1 progeny for analysis. To obtain F3 progeny, 10 P0 worms were plated per seeded plate. These worms laid F1 embryos, which grew to become gravid adults on the same plates (~1000 F1 worms per plate). Five days after plating, the F1 worms on these plates were hypochlorite treated to obtain F2 embryos. Approximately 1000 F2 embryos were plated per new seeded plate, and these worms were hypochlorite treated 3 days later to obtain F3 progeny. In all cases, hypochlorite treatment was performed prior to worms starving on plates.

### L1 starvation survival

Embryos were suspended following hypochlorite treatment in virgin S-basal (no cholesterol or ethanol) at a density of 1 embryo/μL in a volume of 5 mL in a 16 mm glass test tube and placed on a tissue culture roller drum at approximately 25 rpm and 21-22°C. Beginning at day 1 (24 hr after hypochlorite treatment) and continuing every other day, a 100 μL sample was taken and plated on a 6 cm NGM plate with a spot of OP50 in the center. The sample was plated to the side of the OP50. The number of worms plated was scored (total plated = T_P_). Two days later, the number of worms that were alive on the plate were scored (total alive = T_A_). The proportion alive at each time point was calculated as T_A_ / T_P_.

### Brood size

Worms were singled onto 6 cm NGM plates with OP50 as L4 larvae. Worms were transferred to new plates each day until they stopped producing progeny. The number of progeny per plate was scored 2 days after removal of the mother. The total brood size per worm was determined by summing the progeny per plate across all plates for a single worm. Worms were censored if they died during egg laying. This affected 11 worms total in P0 brood size (5 post-dauers, 6 controls). In the F3 generation following long-term dauer, for 0 days of arrest, number of worms that died during egg laying included 0 controls and 1 post-dauer. For 8 days of arrest, number of worms that died during egg laying include 16 controls and 14 post-dauers; 2 controls and 2 post-dauers were sterile. In the F3 generation following short-term dauer, 9 control worms and 9 post-dauer worms died during egg laying; 1 post-dauer worm was sterile.

### Embryo length

Embryos were plated onto unseeded 10 cm NGM plates. Embryos were imaged with a Zeiss Discovery. V20 stereomicroscope with a 10x objective (KSC 190-975). The images were analyzed with FIJI as described previously (44). Lengths of embryos were measured by thresholding embryos and calculating the long axis from ellipse fitting. The background was subtracted, images were thresholded, converted to binary, holes were filled, and particles were analyzed. Analysis was done in batch and the results were manually curated to ensure only quality embryo images were included.

### Worm length

L1 larvae that had been starved for 1 or 8 days were recovered by plating on 10 cm NGM plates with OP50 as previously described (44). After 48 hours of recovery, worms were washed off the plates with virgin S-basal and plated on unseeded 10 cm NGM plates for imaging. Images were taken on a ZeissDiscovery.V20 stereomicroscope with automated zoom. Images were analyzed with the WormSizer plugin for FIJI to determine worm length and manually passed or failed (45).

### Lifespan

For each condition and replicate, 150 L1 larvae arrested for 1 day in virgin S-basal were plated on 6 cm NGM plates seeded with OP50. After two days, 72 worms were randomly picked onto new seeded plates in groups of 12. Adults were picked away from their progeny onto fresh plates every day until egg laying ceased. Worms that responded to gentle prodding with a platinum wire were transferred to fresh plates every 2 - 3 days. Worms were considered dead when they failed to respond to prodding. Worms that crawled off the agar were considered lost and subtracted from the total *n*. No other animals were censored. Lifespan curves were analyzed to determine mean survival using OASIS (46).

### Statistical analysis

Statistics were calculated in R or Microsoft Excel. To test for differences in means across groups, linear mixed effect models were fit to the data using the “nlme” package in R. The summary function was used to calculate p-values for the models, which implements the Wald test. Fixed effects included condition (post-dauer vs. control) and, where applicable, length of starvation (0 or 1 day vs. 8 days). An interaction term was included for experiments with both types of fixed effects. A random effect of biological replicate was included for all models. To test for differences in inter-individual variation across conditions, data were mean normalized within each biological replicate and condition, individual worms across all replicates were pooled, and Levene’s test was used to assess homogeneity of variance across conditions. For starvation survival (Figs 2,4, and 5), logistic curves were fit to survival data and median survival times were calculated (47). Paired t-tests were performed on median survival times. Statistical tests and significance are indicated in figure legends. Plots were generated using ggplot2 in R.

### Sample collection for RNA-seq

F3 embryos were suspended at 1 embryo/μL in S-complete either with or without 25 mg/mL HB101 to obtain fed or starved L1s, respectively. At least 10,000 embryos were used per condition per replicate. Fed larvae were collected 18 hr after hypochlorite treatment as early-stage developing L1 larvae. Starved larvae were collected 24 hr after hypochlorite treatment as arrested L1 larvae that had hatched approximately 12 hr earlier. To collect starved samples, cultures were transferred to 15 mL conical tubes and spun for 1 minute at 3000 rpm. Liquid was aspirated to < 100 uL containing the worm pellet. Worms were washed 0-1X with S-basal. The worm pellet was transferred to a 1.5 mL microcentrifuge tube, flash frozen in liquid nitrogen, and stored at −80°C until RNA isolation. To collect fed samples, cultures were transferred to 15 mL conical tubes and spun at 3000 rpm for 10 seconds. Liquid was aspirated to < 100 uL containing the worm pellet. The pellet was quickly washed 3-4X with 10 mL S-basal, visually inspected to ensure removal of the vast majority of bacteria, transferred to a 1.5 mL microcentrifuge tube, flash frozen, and stored at −80°C until RNA isolation.

### RNA isolation and library preparation

RNA was isolated using TRIzol (Invitrogen) using the manufacturer’s protocol with minor modifications. 1 mL of TRIzol was used per sample along with 100 uL of acid-washed sand. mRNA-seq libraries were prepared using the NEBNext Ultra RNA Library Prep Kit for Illumina (E7530) in two batches, utilizing either 500 or 100 ng of starting RNA per library and 12 or 15 PCR cycles, respectively. Libraries were sequenced using Illumina HiSeq 4000 to obtain single-end 50 bp reads.

### RNA-seq analysis

Bowtie was used to map reads to the WS210 genome (48). We also included transcripts annotated in WS220 mapped back to the WS210 genome coordinates, as described previously (49). Mapping efficiencies ranged from 81-86% for all libraries. HTSeq was used to generate count tables for each library (50). Count tables were analyzed for differential expression using the edgeR package in R, which utilizes a negative binomial model to estimate dispersions (51). Detected genes were considered those expressed at a level of at least 4 counts per million (CPM) across all libraries for all conditions, reducing the number of genes included in the analysis to 8649. The “calcNormFactors” function was used to normalize for RNA composition and the tagwise dispersion estimate was used for differential expression. The exact test was used for pairwise comparisons. Log2 fold change estimates from differential expression analysis in edgeR were used for generating plots in Fig 7 and Supplementary Fig 4. Kolmogorov- Smirnov tests were used to determine differences in cumulative distributions of log2 fold changes.

### Data availability

Raw and processed RNA-seq data is available through the GEO NCBI database with accession number GSE113500.

## SUPPORTING INFORMATION

**Supplementary Fig 1:**
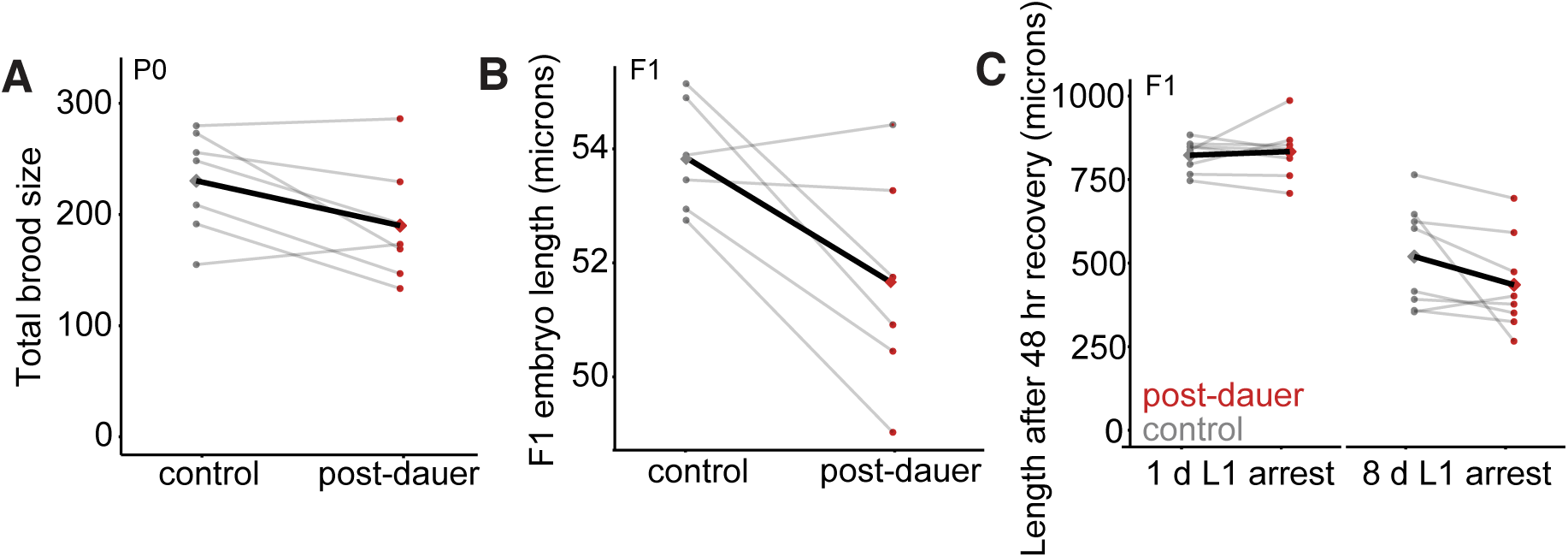
Proximal effects of long--term dauer on brood size, embryo length, and body size. A-C. Each point represents the mean of individual worms for the indicated trait within a biological replicate. Paired replicates are connected with a gray line. Grand means are connected with a black line.

**Supplementary Fig 2:**
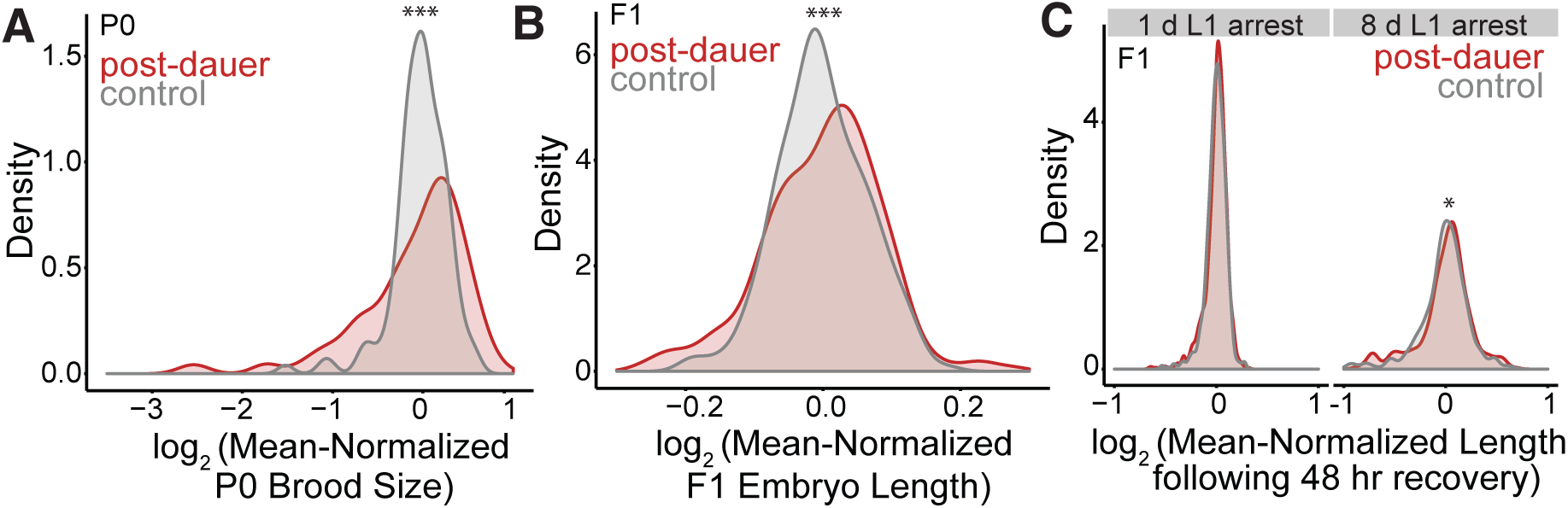
Long-term dauer diapause promotes variability in F1 progeny number and size. A. Brood size data were pooled across 7 biological replicates including 115 post-dauer worms and 120 control worms, p = 2.2 x 10^−8^. B. Embryo length data were pooled across 6 replicates including 256 post-dauer embryos and 235 control embryos, p = 0.00033. C. Worm body lengths following recovery following 1 or 8 days of L1 arrest in F1 progeny of post-dauers and controls were pooled from 8 biological replicates including 748 controls starved for 1 day, 897 post-dauers starved for 1 day, 421 controls starved for 8 days, and 337 post-dauers starved for 8 days. For 1 day, p = 0.13; for 8 days, p = 0.02. A-C. Trait values for individual post-dauer and control worms were mean-normalized to the mean trait value for the corresponding condition and replicate. Data were pooled across biological replicates. Levene’s test was used to assess differences in variance. *p<0.05, ***p<0.001

**Supplementary Fig 3:**
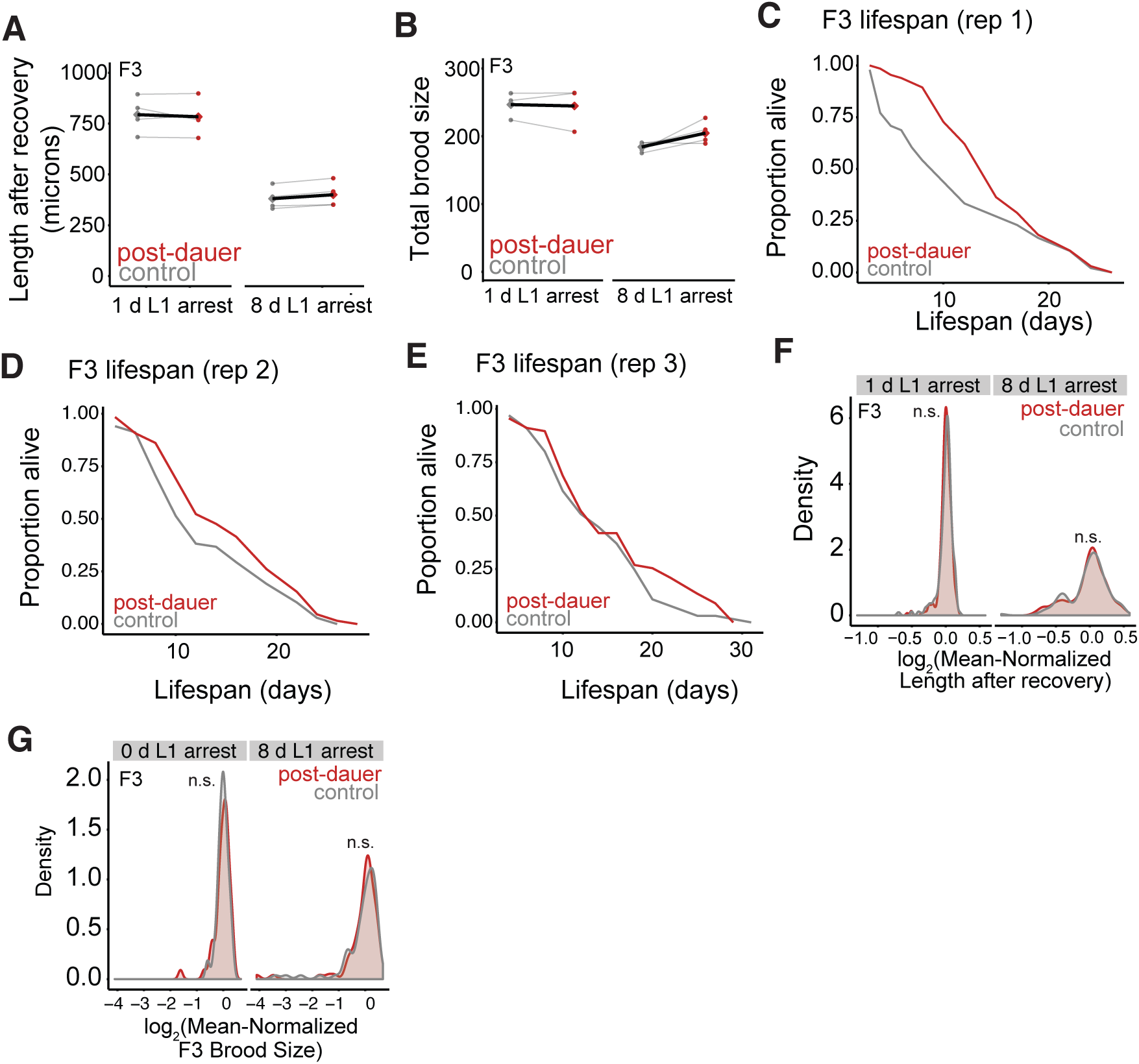
Transgenerational effects of long-term dauer diapause in the F3 generation. A-B. Each point represents the mean of individual worms in the F3 generation for the indicated trait within a biological replicate. Paired replicates are connected with a gray line. Grand means are connected with a black line. C. Replicate 1 of post-dauer and control F3 lifespan. 47 control and 66 post-dauer F3 worms assayed. Control mean: 12.1 days; post-dauer mean: 15 days D. Replicate 2 of post-dauer and control F3 lifespan. 64 control and 64 post-dauer F3 worms assayed. Control mean: 13.6 days; post-dauer mean: 15.5 days E. Replicate 3 of post-dauer and control F3 lifespan. 63 control and 64 post-dauer F3 worms assayed. Control mean: 14.4 days; post-dauer mean: 16.2 days F. Worm body length measurements were pooled across replicates to test for differences in variance (same data as 3B), For 1 day, p = 0.089; for 8 days, p = 0.82 G. Brood sizes data were pooled across replicates to test for differences in variance (same data as Fig 3C). For 0 d: p = 0.18; for 8 d: p = 0.48. F-G. Trait values for individual post--dauer and control worms were mean--normalized to the mean trait value for the corresponding condition and replicate. Data were pooled across biological replicates. Levene’s test was used to assess differences in variance. n.s. not significant

**Supplementary Fig 4:**
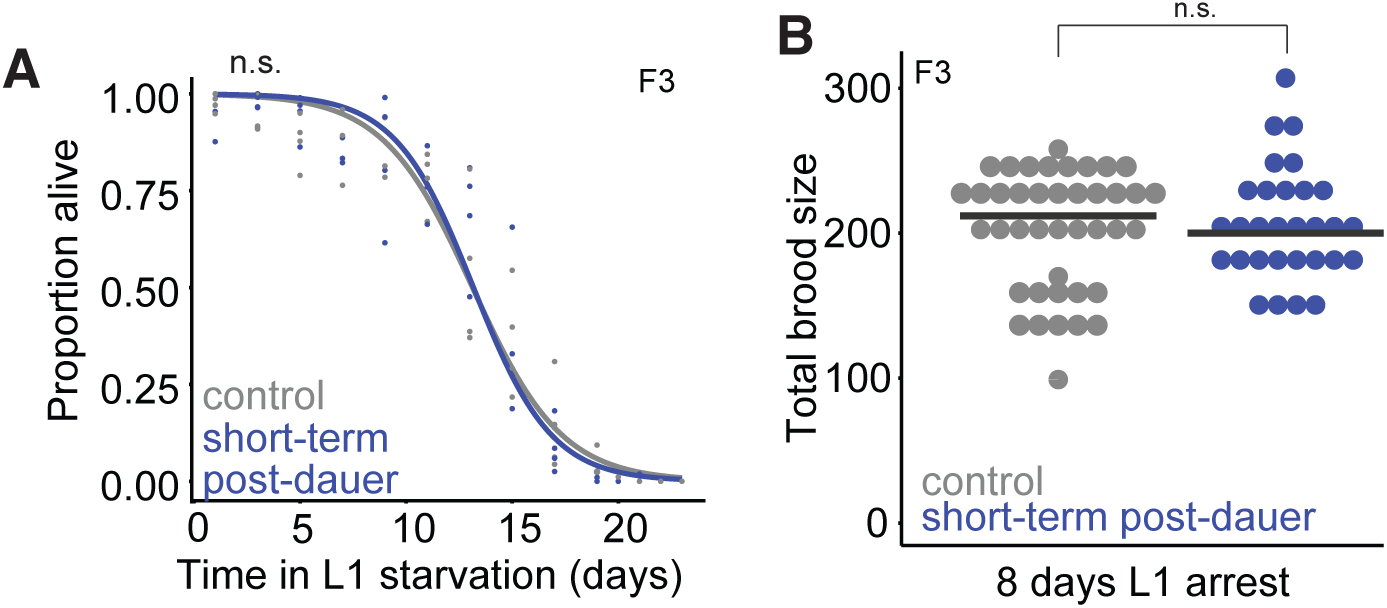
Transgenerational starvation resistance is not apparent in the F3 generation following short-term dauer diapause. A. Starvation survival in the F3 progeny of controls and short-term dauers (6 days as a dauer). 4 biological replicates were scored, logistic curves were fit to data, and median survival times were determined. Paired t-test on median half-lives, p = 0.73. B. Brood size of F3 progeny of controls and short-term dauers was scored after worms experienced 8 days of L1 arrest. 3 biological replicates of 5-18 individual worms per condition were scored. Grand mean for control: 200; for post-dauers: 201. A linear mixed-effect model was fit to brood size data with condition (post-dauer vs. control) as a fixed effect and biological replicate as a random effect. P-values were calculated using the Wald test. Effect of condition, p = 0.75. n.s.: not significant. Horizontal black lines represent median.

**Supplementary Fig 5:**
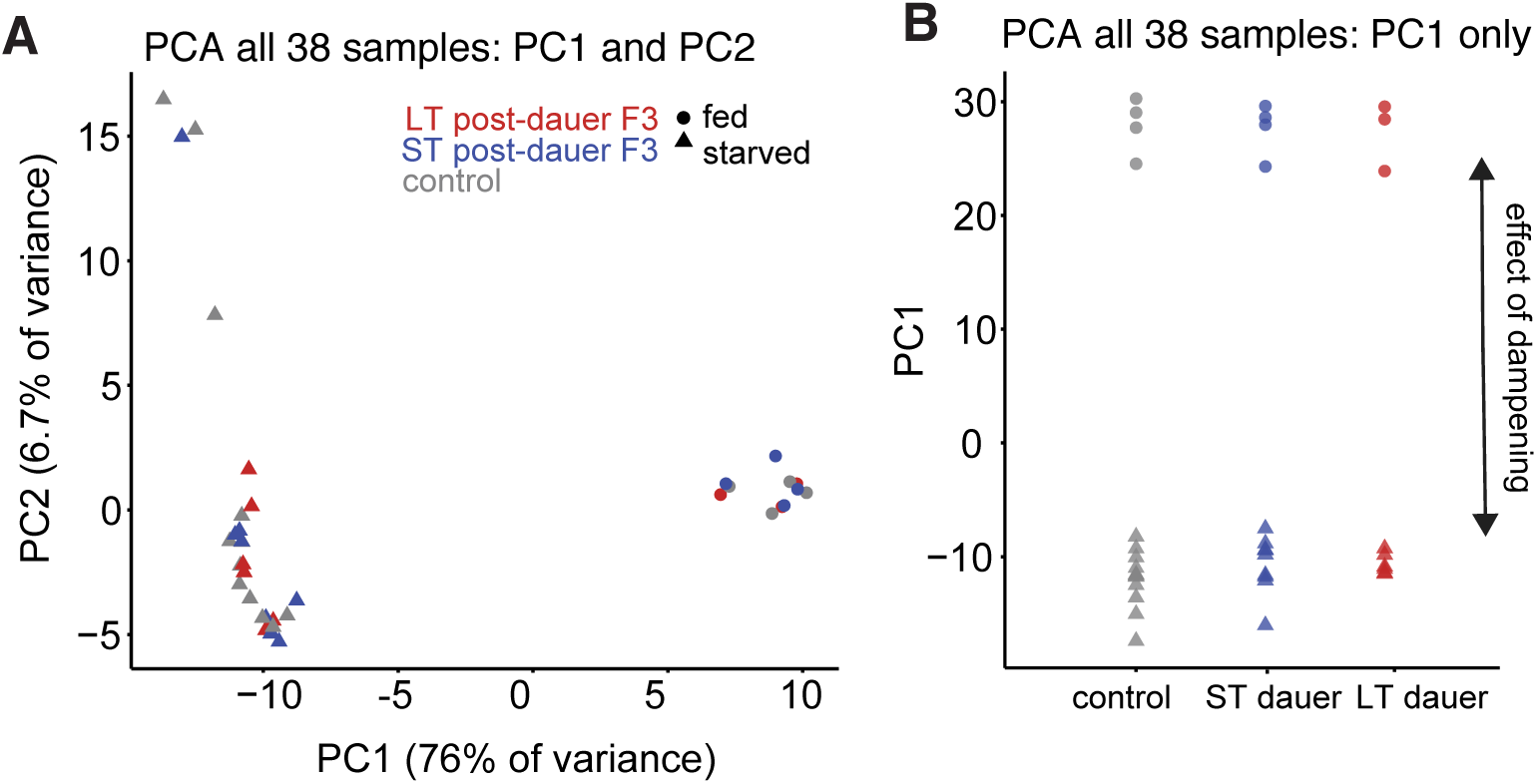
Individual replicates separate on PCA. A. Principal component analysis (PC1 and PC2) of all biological replicates included in differential expression analysis. B. Biological replicates plotted in PC1 only. Considering both current environment (fed vs. starved) and ancestral environment (control, short-term dauer, and long-term dauer) as factors, both effects are significant in a multivariate ANOVA using Wilks’ Lambda for PC1. Effect of current environment, p < 2.2 x 10^−16^; effect of ancestral environment, p = 0.00022.

**Supplementary Fig 6:**
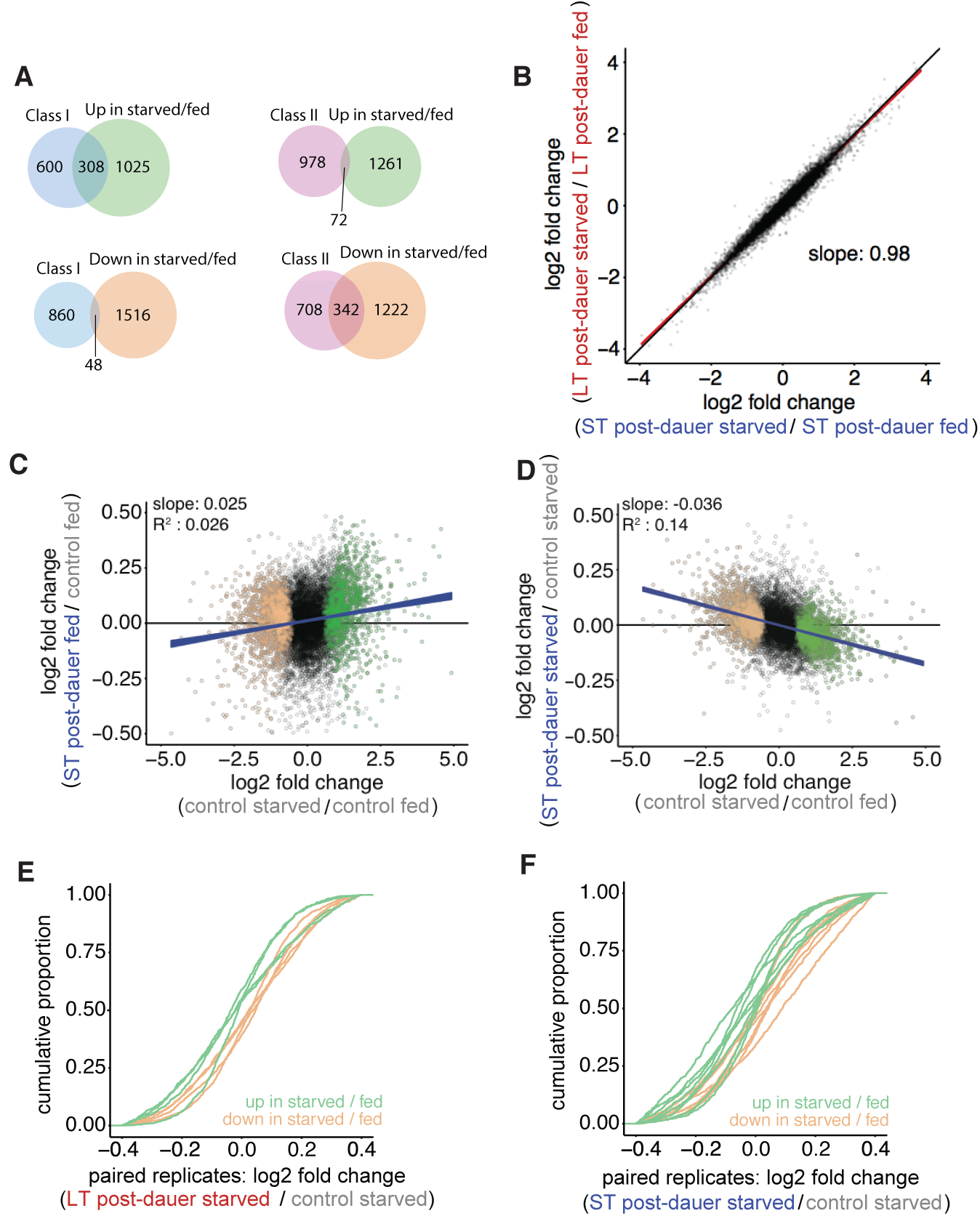
Additional RNA-seq analysis. A. Class I and Class II targets defined as genes positively or negatively regulated by DAF--16. Class I and Class II targets were filtered to include those with FDR < 0.05 that were also in the background set of 8649 genes included in starved / fed differential expression analysis. Class I targets are over-enriched in genes up in starved / fed (hypergeometric p = 1 x 10^−49^). Class II targets are over-enriched among genes in starved / fed (hypergeometric p = 2.6 x 10^−34^). Class I targets can explain 23.1% of starvation upregulation; class II targets can explain 21.9% of starvation downregulation. B. Log2 fold change of all genes in ST post-dauer starved / ST post-dauer fed compared to log2 fold change in LT post-dauer starved / LT post-dauer fed. Red line indicates simple linear regression through all points, and thickness of line indicates 95% confidence interval. Slope is 0.98, with confidence interval ranging from 0.973 to 0.981. C-D. Genes up in control starved / control fed using FDR < 1 x 10^−10^ are plotted in green. Genes down in control starved / control fed are plotted in tan. Blue line is a simple linear regression through all points, and thickness of line indicates confidence interval. C. Log2 fold changes of all genes in control starved / control fed plotted against log2 fold changes of ST post-dauer fed F3 / control fed plotted in black. D. Log2 fold changes of all genes in control starved / control fed plotted against log2 fold changes of ST post-dauer starved F3 / control starved plotted in black. E-F. Cumulative distribution function plots for individual paired replicates using the same groups of genes defined as up- and down-regulated in control starved / control fed comparison. Paired t-test on median fold changes for these two gene groups. E. LT post-dauer starved / control starved comparison, p = 0.0083. F. ST post-dauer starved / control starved comparison, p = 0.088.

**Supplementary Table 1:** Starvation survival and lifespan statistics.

| <u>Starvation survival</u> |  |  |  |  |  |  |
| --- | --- | --- | --- | --- | --- | --- |
| **F1 progeny of long-term dauer and control** | **Avg median survival (days)** | **StDev (days)** | **n** | **Avg delta compared to control (days)** | **StDev of deltas compared to control (days)** | **p-value vs. control** |
| Control | 11 | 3 | 7 | N/A | N/A | N/A |
| Long-term dauer | 9.1 | 3.1 | 7 | -1.9 | 2.6 | 0.11 |
| **F3 progeny of long-term dauer and control** | **Avg median survival (days)** | **StDev (days)** | **n** | **Avg delta compared to control (days)** | **StDev of deltas compared to control (days)** | **p-value vs. control** |
| Control | 10.5 | 1.1 | 8 | N/A | N/A | N/A |
| Long-term dauer | 11.9 | 0.6 | 8 | 1.4 | 1.1 | 0.01 |
| **F3 progeny of short-term dauer and control** | **Avg median survival (days)** | **StDev (days)** | **n** | **Avg delta compared to control (days)** | **StDev of deltas compared to control (days)** | **p-value vs. control** |
| Control | 13.3 | 1.6 | 4 | N/A | N/A | N/A |
| Short-term dauer | 13.3 | 1.5 | 4 | 0.08 | 0.4 | 0.73 |
| <u>Lifespan</u> |  |  |  |  |  |  |
| **F3 progeny of long-term dauer and control** | **Avg median lifespan (days)** | **StDev (days)** | **n** | **Avg delta compared to control (days)** | **StDev of deltas compared to control (days)** | **p-value vs. control** |
| Control | 13.4 | 1.2 | 3 | N/A | N/A | N/A |
| Long-term dauer | 15.6 | 0.6 | 3 | 2.2 | 0.6 | 0.02 |

**Supplementary File 1: edgeR output for pairwise RNA--seq comparisons (.xlsx file)**

## SUPPLEMENTARY DISCUSSION

### Increased phenotypic variation after long-term dauer diapause

In addition to long-term post-dauer worms producing fewer, smaller, starvation-sensitive F1 progeny, there is also greater inter-individual variation in P0 brood size and F1 progeny size. Greater inter-individual variability could occur as a result of 1) decanalization or 2) an evolutionary bet-hedging strategy (52). Decanalization occurs when phenotypic robustness is compromised, as a consequence of, for example, conditions that result in pathology. We believe the observed decreases in average P0 brood size and F1 progeny size and starvation resistance following long-term dauer arrest reflect pathological consequences of extended starvation exceeding the buffering capacity of the organism. Consistent with this interpretation, appreciable lethality and developmental abnormalities affecting the reproductive system have been reported after 60 and 25 days of dauer arrest, respectively (31, 32). Alternatively, bet hedging is thought to be evolutionarily adaptive in unpredictable environments (53). In a bet-hedging scenario, an isogenic population exhibits inter-individual variability such that it is suited for variable environmental conditions. In a particular condition, the bet-hedging population may be less fit than a highly specialized population. However, the bet-hedging population reduces variation in fitness across environments. If post-dauer worms display a bet-hedging strategy, then a subpopulation of post-dauers would be expected to perform better than controls in at least some environmental condition. Our data do not provide compelling evidence for a sub-population of post-dauers or their progeny that performs better than controls for any of the proximal traits assayed, despite increased variability. However, we cannot formally exclude the possibility that there are other environmental conditions that would reveal an advantage to increased phenotypic variation as a consequence of extended dauer diapause. We also emphasize that this consideration of bet hedging is in the context of proximal traits. Since we found that the great-grandprogeny of long-term dauers survive starvation better, it is possible that any advantage to a bet-hedging strategy is only apparent on the scale of multiple generations.

## REFERENCES

1. Kelly SA, Panhuis TM, Stoehr AM. Phenotypic plasticity: molecular mechanisms and adaptive significance. Compr Physiol. 2012;2(2):1417–39.

2. Rapp RA, Wendel JF. Epigenetics and plant evolution. New Phytol. 2005;168(1):81–91.

3. Heard E, Martienssen RA. Transgenerational epigenetic inheritance: myths and mechanisms. Cell. 2014;157(1):95–109.

4. Jablonka E, Raz G. Transgenerational epigenetic inheritance: prevalence, mechanisms, and implications for the study of heredity and evolution. Q Rev Biol. 2009;84(2):131–76.

5. Greer EL, Maures TJ, Ucar D, Hauswirth AG, Mancini E, Lim JP, et al. Transgenerational epigenetic inheritance of longevity in Caenorhabditis elegans. Nature. 2011;479(7373):365–71.

6. Buckley BA, Burkhart KB, Gu SG, Spracklin G, Kershner A, Fritz H, et al. A nuclear Argonaute promotes multigenerational epigenetic inheritance and germline immortality. Nature. 2012;489(7416):447–51.

7. Schott D, Yanai I, Hunter CP. Natural RNA interference directs a heritable response to the environment. Sci Rep. 2014;4:7387.

8. Ni JZ, Kalinava N, Chen E, Huang A, Trinh T, Gu SG. A transgenerational role of the germline nuclear RNAi pathway in repressing heat stress-induced transcriptional activation in C. elegans. Epigenetics Chromatin. 2016;9:3.

9. Minkina O, Hunter CP. Stable Heritable Germline Silencing Directs Somatic Silencing at an Endogenous Locus. Molecular cell. 2017;65(4):659–70e5.

10. Klosin A, Casas E, Hidalgo-Carcedo C, Vavouri T, Lehner B. Transgenerational transmission of environmental information in C. elegans. Science. 2017;356(6335):320–3.

11. Schindler AJ, Baugh LR, Sherwood DR. Identification of late larval stage developmental checkpoints in Caenorhabditis elegans regulated by insulin/IGF and steroid hormone signaling pathways. PLoS Genet. 2014;10(6):e1004426.

12. Baugh LR. To grow or not to grow: nutritional control of development during Caenorhabditis elegans L1 arrest. Genetics. 2013;194(3):539–55.

13. Angelo G, Van Gilst MR. Starvation protects germline stem cells and extends reproductive longevity in C. elegans. Science. 2009;326(5955):954–8.

14. Schulenburg H, Felix MA. The Natural Biotic Environment of Caenorhabditis elegans. Genetics. 2017;206(1):55-86.

15. Hu PJ. Dauer. WormBook. 2007:1-19.

16. Kostal V. Eco-physiological phases of insect diapause. J Insect Physiol. 2006;52(2):113–27.

17. Wang J, Kim SK. Global analysis of dauer gene expression in Caenorhabditis elegans. Development. 2003;130(8):1621–34.

18. Baugh LR, Kurhanewicz N, Sternberg PW. Sensitive and precise quantification of insulin-like mRNA expression in Caenorhabditis elegans. PLoS One. 2011;6(3):e18086.

19. McElwee JJ, Schuster E, Blanc E, Thomas JH, Gems D. Shared transcriptional signature in Caenorhabditis elegans Dauer larvae and long-lived daf-2 mutants implicates detoxification system in longevity assurance. J Biol Chem. 2004;279(43):44533–43.

20. Fielenbach N, Antebi A. C. elegans dauer formation and the molecular basis of plasticity. Genes Dev. 2008;22(16):2149–65.

21. Li W, Kennedy SG, G. Ruvkun daf-28 encodes a C. elegans insulin superfamily member that is regulated by environmental cues and acts in the DAF-2 signaling pathway. Genes Dev. 2003;17(7):844–58.

22. Ren P, Lim CS, Johnsen R, Albert PS, Pilgrim D, Riddle DL. Control of C. elegans larval development by neuronal expression of a TGF-beta homolog. Science. 1996;274(5291):1389–91.

23. Schackwitz WS, Inoue T, Thomas JH. Chemosensory neurons function in parallel to mediate a pheromone response in C. elegans. Neuron. 1996;17(4):719–28.

24. Bargmann CI. Chemosensation in C. elegans. WormBook. 2006:1–29.

25. Dixon SJ, Alexander M, Chan KK, Roy PJ. Insulin-like signaling negatively regulates muscle arm extension through DAF-12 in Caenorhabditis elegans. Dev Biol. 2008;318(1):153–61.

26. Albert PS, Riddle DL. Developmental alterations in sensory neuroanatomy of the Caenorhabditis elegans dauer larva. J Comp Neurol. 1983;219(4):461–81.

27. Keane J, Avery L. Mechanosensory inputs influence Caenorhabditis elegans pharyngeal activity via ivermectin sensitivity genes. Genetics. 2003;164(1):153–62.

28. Cunningham KA, Ashrafi K. Fat rationing in dauer times. Cell Metab. 2009;9(2):113–4.

29. Narbonne P, Roy R. Caenorhabditis elegans dauers need LKB1/AMPK to ration lipid reserves and ensure long-term survival. Nature. 2009;457(7226):210–4.

30. Barriere A, Felix MA. High local genetic diversity and low outcrossing rate in Caenorhabditis elegans natural populations. Curr Biol. 2005;15(13):1176–84.

31. Klass M, Hirsh D. Non-ageing developmental variant of Caenorhabditis elegans. Nature. 1976;260(5551):523–5.

32. Kim S, Paik YK. Developmental and reproductive consequences of prolonged non-aging dauer in Caenorhabditis elegans. Biochem Biophys Res Commun. 2008;368(3):588–92.

33. Skinner MK. What is an epigenetic transgenerational phenotype? F3 or F2. Reprod Toxicol. 2008;25(1):2–6.

34. Tepper RG, Ashraf J, Kaletsky R, Kleemann G, Murphy CT, Bussemaker HJ. PQM-1 complements DAF-16 as a key transcriptional regulator of DAF-2-mediated development and longevity. Cell. 2013;154(3):676–90.

35. Chen J, Nolte V, Schlotterer C. Temperature stress mediates decanalization and dominance of gene expression in Drosophila melanogaster. PLoS Genet. 2015;11(2):e1004883.

36. Jobson MA, Jordan JM, Sandrof MA, Hibshman JD, Lennox AL, Baugh LR. Transgenerational Effects of Early Life Starvation on Growth, Reproduction, and Stress Resistance in Caenorhabditis elegans. Genetics. 2015;201(1):201–12.

37. Hall SE, Beverly M, Russ C, Nusbaum C, Sengupta P. A cellular memory of developmental history generates phenotypic diversity in C. elegans. Curr Biol. 2010;20(2):149–55.

38. Calabrese EJ, Baldwin LA. Defining hormesis. Hum Exp Toxicol. 2002;21(2):91–7.

39. Rechavi O, Houri-Ze’evi L, Anava S, Goh WS, Kerk SY, Hannon GJ, et al. Starvation-induced transgenerational inheritance of small RNAs in C. elegans. Cell. 2014;158(2):277–87.

40. Murren CJ, Auld JR, Callahan H, Ghalambor CK, Handelsman CA, Heskel MA, et al. Constraints on the evolution of phenotypic plasticity: limits and costs of phenotype and plasticity. Heredity (Edinb). 2015;115(4):293–301.

41. Dewitt TJ, Sih A, Wilson DS. Costs and limits of phenotypic plasticity. Trends Ecol Evol. 1998;13(2):77–81.

42. Herman JJ, Spencer HG, Donohue K, Sultan SE. How stable ‘should’ epigenetic modifications be? Insights from adaptive plasticity and bet hedging. Evolution. 2014;68(3):632–43.

43. Felix MA, Braendle C. The natural history of Caenorhabditis elegans. Curr Biol. 2010;20(22):R965–9.

44. Hibshman JD, Hung A, Baugh LR. Maternal Diet and Insulin-Like Signaling Control Intergenerational Plasticity of Progeny Size and Starvation Resistance. PLoS Genet. 2016;12(10):e1006396.

45. Moore BT, Jordan JM, Baugh LR. WormSizer: high-throughput analysis of nematode size and shape. PLoS One. 2013;8(2):e57142.

46. Yang JS, Nam HJ, Seo M, Han SK, Choi Y, Nam HG, et al. online application for the survival analysis of lifespan assays performed in aging research. PLoS One. 2011;6(8):e23525.

47. Kaplan RE, Chen Y, Moore BT, Jordan JM, Maxwell CS, Schindler AJ, et al. dbl-1/TGF-beta and daf-12/NHR Signaling Mediate Cell-Nonautonomous Effects of daf-16/FOXO on Starvation-Induced Developmental Arrest. PLoS Genet. 2015;11(12):e1005731.

48. Langmead B, Trapnell C, Pop M, Salzberg SL. Ultrafast and memory-efficient alignment of short DNA sequences to the human genome. Genome Biol. 2009;10(3):R25.

49. Maxwell CS, Antoshechkin I, Kurhanewicz N, Belsky JA, Baugh LR. Nutritional control of mRNA isoform expression during developmental arrest and recovery in C. elegans. Genome Res. 2012;22(10):1920–9.

50. Anders S, Pyl PT, Huber W. HTSeq--a Python framework to work with high-throughput sequencing data. Bioinformatics. 2015;31(2):166–9.

51. Robinson MD, McCarthy DJ, Smyth GK. edgeR: a Bioconductor package for differential expression analysis of digital gene expression data. Bioinformatics. 2010;26(1):139–40.

52. Waddington CH. The strategy of the genes; a discussion of some aspects of theoretical biology. London,: Allen & Unwin; 1957. ix, 262 p. p.

53. Starrfelt J, Kokko H. Bet-hedging--a triple trade-off between means, variances and correlations. Biol Rev Camb Philos Soc. 2012;87(3):742–55.

